# FDA-Approved MEK1/2 Inhibitor, Trametinib, Protects Mice from Cisplatin and Noise-Induced Hearing Loss

**DOI:** 10.1101/2024.05.20.595056

**Authors:** Richard D. Lutze, Matthew A. Ingersoll, Regina G. Kelmann, Tal Teitz

**Affiliations:** Department of Pharmacology and Neuroscience, School of Medicine, Creighton University, Omaha, NE 68178, USA

**Keywords:** MAPK pathway, MEK1/2, trametinib, hearing protection, drug repurposing, oral delivery, cisplatin-induced hearing loss, noise-induced hearing loss

## Abstract

Hearing loss is one of the most common types of disability; however, there is only one FDA-approved drug to prevent any type of hearing loss. Treatment with the highly effective chemotherapy agent, cisplatin, and exposure to high decibel noises are two of the most common causes of hearing loss. The mitogen activated protein kinase (MAPK) pathway, a phosphorylation cascade consisting of RAF, MEK1/2, and ERK1/2, has been implicated in both types of hearing loss. Pharmacologically inhibiting BRAF or ERK1/2 is protective from noise and cisplatin-induced hearing loss in multiple mouse models. Trametinib, a MEK1/2 inhibitor, protects from cisplatin induced outer hair cell death in mouse cochlear explants; however, to the best of our knowledge, inhibiting MEK1/2 has not yet been shown to be protective from hearing loss *in vivo*. In this study, we demonstrate that trametinib protects from cisplatin-induced hearing loss in a translationally relevant mouse model and does not interfere with cisplatin’s tumor killing efficacy in cancer cell lines. Higher doses of trametinib were toxic to mice when combined with cisplatin but lower doses of the drug were protective from hearing loss without any known toxicity. Trametinib also protected mice from noise-induced hearing loss and synaptic damage. This study shows that MEK1/2 inhibition protects from both insults of hearing loss and that targeting all three kinases in the MAPK pathway protect from cisplatin and noise-induced hearing loss in mice.

## Introduction

Approximately 20% of the world population is estimated to have hearing loss with over 400 million individuals having moderate to severe hearing loss [1,2]. Two of the most common causes of hearing loss are exposure to loud noise and cisplatin administration for the treatment of many types of cancers [3–6]. Hearing loss impedes the development of children and can contribute to dementia and cognitive decline in the elderly [7–10]. There is only one Food and Drug Administration (FDA)-approved drug to prevent any type of hearing loss [11–13], cochlear implants only work in a subset of individuals [14], and hearing aids have low retention rates with people who need them [15]. Therefore, there is a dire need to develop drugs that can prevent hearing loss. Currently, there are many promising therapeutic compounds in preclinical studies and clinical trials that are targeting pathways known to be involved in hearing loss [16–19].

One of these pathways that are known to contribute to hearing loss is the mitogen-activated protein kinase (MAPK) pathway. Many laboratories, including our own, have demonstrated that this pathway is activated immediately after noise exposure and cisplatin administration with inhibition of this pathway through genetic manipulation, or pharmacological agents leading to protection from hearing loss [20–29]. The MAPK pathway consists of three main kinases, RAF, MEK1/2, and ERK1/2 (Figure 1A) [30]. Activation of this phosphorylation cascade leads to a wide range of cellular responses such as proliferation, cell migration, cell survival, inflammation, and apoptosis [31–33]. The cellular response that occurs following the activation of the MAPK pathway is typically dependent on the type of stimulus, duration of stimulus, and the cell type [24,33]. Overactivation of MAPK proteins has been studied extensively as a main cause of cancer progression; however, this pathway is implicated in hearing loss and inhibition of the pathway leads to significant hearing protection in many different strains of mice [24–27,31].

**Figure 1:**
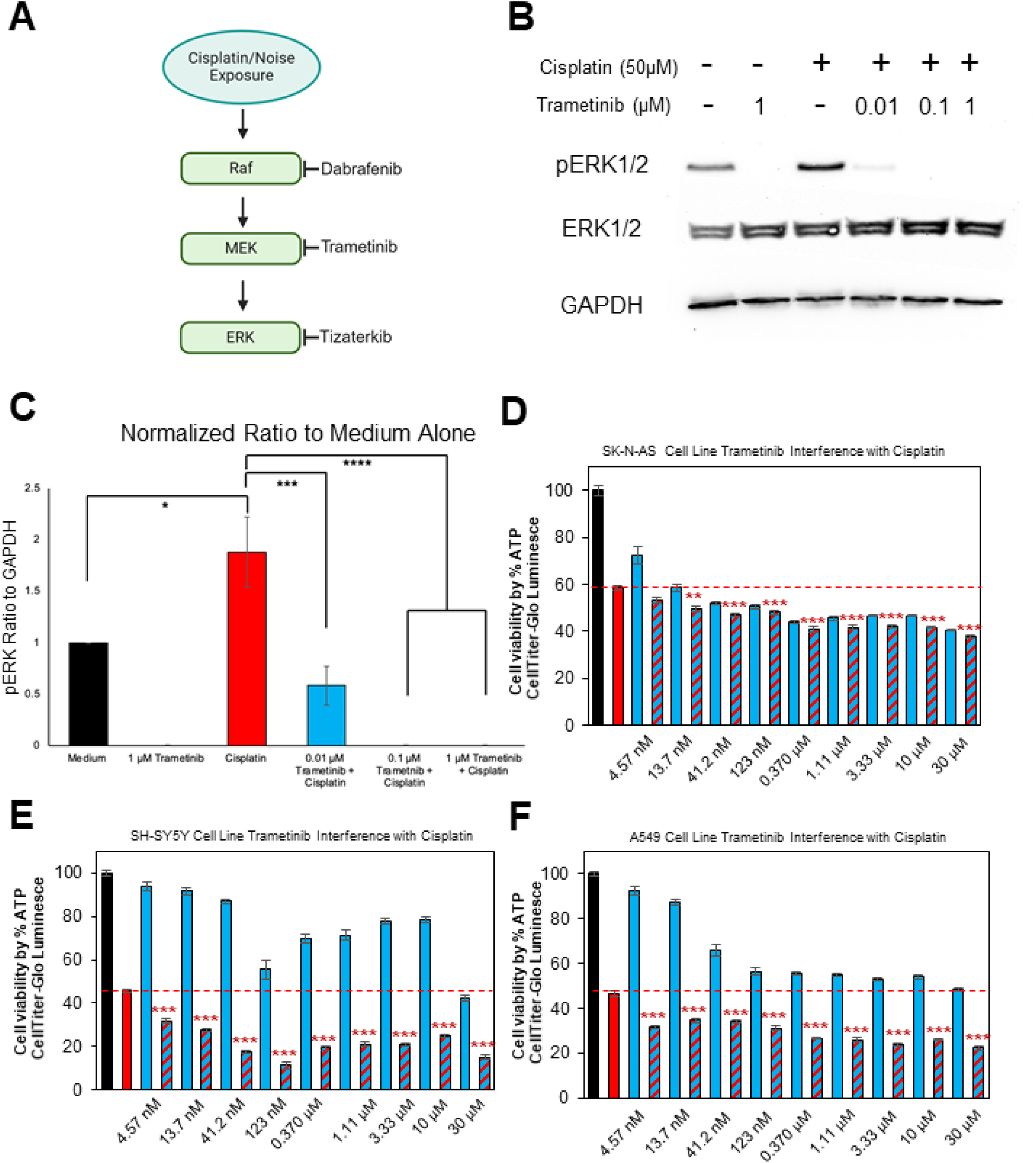
Trametinib inhibits MAPK activity in the HEI-OC1 cell line and does not interfere with cisplatin’s tumor killing ability in cancer cell lines. **(A)** Schematic of the MAPK phosphorylation cascade in which dabrafenib inhibits BRAF, trametinib inhibits MEK1/2, and tizaterkib inhibits ERK1/2. **(B)** Representative western blots of HEI-OC1 cell lysates treated with medium, cisplatin, and different concentrations of trametinib. Treatment groups from left to right are as follows: medium alone, 1μM trametinib alone, 50μM cisplatin alone, 50μM cisplatin + 0.01μM trametinib, 50μM cisplatin + 0.1μM trametinib, and 50μM cisplatin + 1μM trametinib. **(C)** Quantification of western blots represented in (A). A total of 4 separate experiments were performed. The ratio of pERK to GAPDH was measured for each individual lane and all groups were then normalized to the medium alone treatment group. Data shown as means ± SEM. *P<0.05, ***P<0.001, ****P<0.0001. All groups compared to one another by one-way ANOVA with Bonferroni post hoc test. **(D)** Percentage of cell viability for SK-N-AS cell line treated with cisplatin and various concentrations of trametinib starting at 4.57nM and going up to 30µM in increments of 3 fold increases. **(E)** Percentage of cell viability for SH-SY5Y cell line treated with cisplatin and various concentrations of trametinib as mentioned in (D). **(F)** Percentage of cell viability for A549 cell line treated with cisplatin and various concentrations of trametinib as mentioned in (D). Medium alone (Black), Cisplatin alone (Red), trametinib alone (Blue), and trametinib + cisplatin (blue and red checkered pattern). All wells treated with cisplatin had the same concentration of cisplatin and increasing concentrations of trametinib were used starting at 4.57nM going up to 30µM going from left to right. All treatments were normalized to medium alone treated cells and compared to the cisplatin alone treatment. Data shown as mean ± SEM *P<0.05, **P<0.01, ***P<0.001 compared to cisplatin alone by one-way ANOVA with Bonferroni post hoc test.

Our laboratory has demonstrated that dabrafenib, an FDA-approved BRAF inhibitor, protects mice from both cisplatin and noise-induced hearing loss (NIHL) [24,25]. Additionally, we have shown that tizaterkib, an ERK1/2 inhibitor that is in phase 1 clinical trials for the treatment of solid tumors and hematological malignancies, protects mice from NIHL at low doses [27]. To the best of our knowledge, MEK 1/2 inhibition has not been shown to protect from hearing loss *in vivo*. Inhibition of upstream BRAF and downstream ERK1/2 protects from hearing loss; however, demonstrating that MEK1/2 inhibition protects from hearing loss is needed to confirm that the canonical MAPK pathway of BRAF, MEK1/2, and ERK1/2 contributes to hearing loss [24–27].

Trametinib is a highly selective MEK1/2 inhibitor that is FDA-approved for the treatment of melanoma, non-small cell lung cancer, thyroid cancer, solid tumors, and low-grade glioma with BRAF V600 mutations [34–37]. Trametinib is used in combination with dabrafenib for the treatment of many different cancers and is well-tolerated by many patients [34]. Trametinib has many promising properties to be repurposed as an otoprotective agent. FDA-approved drugs take much less time to get to market for the treatment of another indication and is much cheaper than getting new chemical entities FDA-approved [38]. As previously mentioned, trametinib is highly selective for MEK1/2 with an IC_50_ lower than 10nM for the respective kinases [39,40]. This high selectivity allows for lower doses of the drug to be administered and limits off target side effects. Additionally, dabrafenib has been shown to be protective from hearing loss, which inhibits upstream BRAF [24,25]. It is conceivable that targeting MEK1/2 could be more efficacious in protecting from hearing loss because it is further down in the MAPK pathway than BRAF (Figure 1A). Targeting kinases more downstream in the MAPK pathway can have greater modulation of the pathway than upstream inhibitors which could end in better hearing outcomes after noise exposure or cisplatin administration [30,41].

In this study, we tested the ability of trametinib to protect from both cisplatin and noise-induced hearing loss. Previously, trametinib was shown to protect cochlear explants from cisplatin-induced outer hair cell death with an IC_50_ of 100nM [25]. Trametinib’s efficacy in protecting from hearing loss *in vivo* is unknown. We first demonstrated that trametinib inhibits the MAPK pathway and does not interfere with cisplatin’s tumor killing efficacy in cancer cell lines. We then tested the ability of trametinib to protect from cisplatin-induced hearing loss by utilizing a clinically, relevant mouse model of cisplatin ototoxicity [42–44]. Two different functional hearing tests, auditory brainstem response (ABR) and distortion product otoacoustic emission (DPOAE) [45,46], were performed to measure mice hearing ability, as well as outer hair cell counts. Finally, we determined whether trametinib protects from NIHL and synaptic damage. This study further supports that the MAPK pathway is a main cellular pathway that causes hearing loss and that pharmacological inhibition of this pathway is a useful therapeutic strategy to protect individuals from both causes of hearing loss.

## Results

### Trametinib inhibits MAPK activation in the HEI-OC1 cell line and does not interfere with cisplatin’s tumor killing ability in multiple cancer cell lines

To confirm that trametinib inhibits MEK1/2, we performed western blots using HEI-OC1 lysates that were treated with cisplatin and different concentrations of trametinib. Six different treatments were performed a total of 4 separate times and 4 individual western blots were run. The six treatment groups are as follows: medium alone, 1μM trametinib alone, 50μM cisplatin alone, 50μM cisplatin + 0.01μM trametinib, 50μM cisplatin + 0.1μM trametinib, and 50μM cisplatin + 1μM trametinib. Phosphorylated ERK1/2 (pERK) was chosen as the protein of interest because ERK1/2 is directly downstream of MEK1/2 in the MAPK pathway and trametinib does not inhibit MEK1/2 phosphorylation (Figure 1A), but it prevents MEK1/2 from having the catalytic ability to activate downstream proteins [40,47]. GAPDH was used as the loading control and the band intensity of pERK was divided by the intensity of GAPDH to get the normalized ratio. All groups were then normalized to the medium alone treatment group. A dose of 50μM cisplatin significantly increased pERK in the HEI-OC1 cell line compared to medium alone and all concentrations of trametinib decreased pERK. There was over a threefold decrease in ERK1/2 phosphorylation following co-administration of 0.01μM trametinib and cisplatin compared to the cisplatin alone treated HEI-OC1 cells. Concentrations of 0.1 and 1μM trametinib completely abrogated ERK1/2 phosphorylation in the presence or absence of cisplatin (Figure 1B & C).

Trametinib was then treated with cisplatin in several different cancer cell lines that cisplatin is commonly used for treatment of the respective tumors (neuroblastoma and lung carcinoma) [48,49]. The CellTiter-Glo Assay was performed to measure the cell viability and determine whether trametinib interferes with cisplatin’s tumor killing ability *in vitro.* In a 96 well plate, 9600 cells were plated into each well and the medium alone wells were considered 100% cell viability. We chose a cisplatin concentration that decreased cell viability by approximately 50% and all wells treated with cisplatin had the same cisplatin concentration. For the wells treated with trametinib, cells were treated with the drug by itself or combined with cisplatin. A wide range of doses were utilized starting at 30µM and serial dilutions of 1/3 were performed until a low dose of 4.57nM was achieved to show that neither high nor low concentrations of trametinib interfere with cisplatin. Cells co-treated with cisplatin and trametinib were compared to cisplatin alone treated wells to determine whether any interference occurred between the two drugs. Each treatment group had 6 wells. For all three cell lines that were treated with both drugs, SK-N-AS (neuroblastoma), SH-SY5Y (neuroblastoma), and A549 (small-cell lung carcinoma), trametinib did not interfere with cisplatin’s tumor killing ability at any of the tested doses (Figure 1D-F). Trametinib and cisplatin treated cells had less cell viability than the cisplatin alone cells, which indicates that trametinib treatment with cisplatin enhances the killing of the tumor cells compared to cisplatin alone. Trametinib by itself reduced cell viability in all 3 cancer cell lines, especially at concentrations of 123nM and higher (Figure 1D-F).

### Trametinib protects from cisplatin-induced hearing loss in a clinically relevant mouse model

To determine whether oral administration of trametinib protects from cisplatin-induced hearing loss, a previously optimized, clinically relevant mouse model of cisplatin administration was utilized [42–44]. Mice were treated with 3mg/kg cisplatin in the morning and treated with trametinib 45 minutes before cisplatin in the morning and again in the evening. There was a total of 4 days of cisplatin treatment and 5 days of trametinib treatment which was followed up with a 9-day recovery period in which no drugs were administered to the mice. This cycle was repeated for a total of 3 times and hearing tests were performed before and after treatment to determine the amount of hearing loss per experimental cohort (Figure 2A). Mice co-treated with 1mg/kg trametinib and cisplatin had significantly lower ABR threshold shifts at 8, 16, and 32 kHz with average threshold shift reductions of 35, 40, and 41 dB compared to the cisplatin alone treated mice, respectively (Figure 2B). Mice co-treated with 0.2mg/kg trametinib and cisplatin had significantly lower ABR threshold shifts of 22 dB at 16 kHz and 24 dB at 32 kHz compared to cisplatin alone treated mice (Figure 2B). 0.1mg.kg trametinib did not significantly decrease ABR threshold shifts compared to cisplatin alone treated mice (Figure 2B). 1mg/kg trametinib co-treated mice with cisplatin had significantly higher ABR wave 1 amplitude at the 16 kHz region compared to cisplatin alone treated mice at 90-, 80-, 70-, and 60 dB while the mice co-treated with 0.2mg/kg trametinib and cisplatin had significantly higher wave 1 amplitudes at 90- and 80 dB (Figure 2C). DPOAE threshold shifts were also measured and 1mg/kg trametinib co-treated mice had significantly lower DPOAE threshold shifts at 12 and 16 kHz compared to the cisplatin alone treatment while 0.2mg/kg trametinib co-treated mice with cisplatin had significantly lower DPOAE threshold shifts at 12 kHz (Figure 2D). Cisplatin alone treated mice had an average DPOAE threshold shift of 40 dB ± 4 and 43 dB ± 4 at the 12 and 16 kHz, respectively, while mice co-treated with 1mg/kg trametinib and cisplatin had an average DPOAE threshold shift of 12 dB ± 4 at 12 kHz and 8 dB ± 6 at 16 kHz. Mice co-treated with 0.2mg/kg trametinib and cisplatin had an average DPOAE threshold shift of 24 dB ± 4 at 12 kHz (Figure 2D).

**Figure 2:**
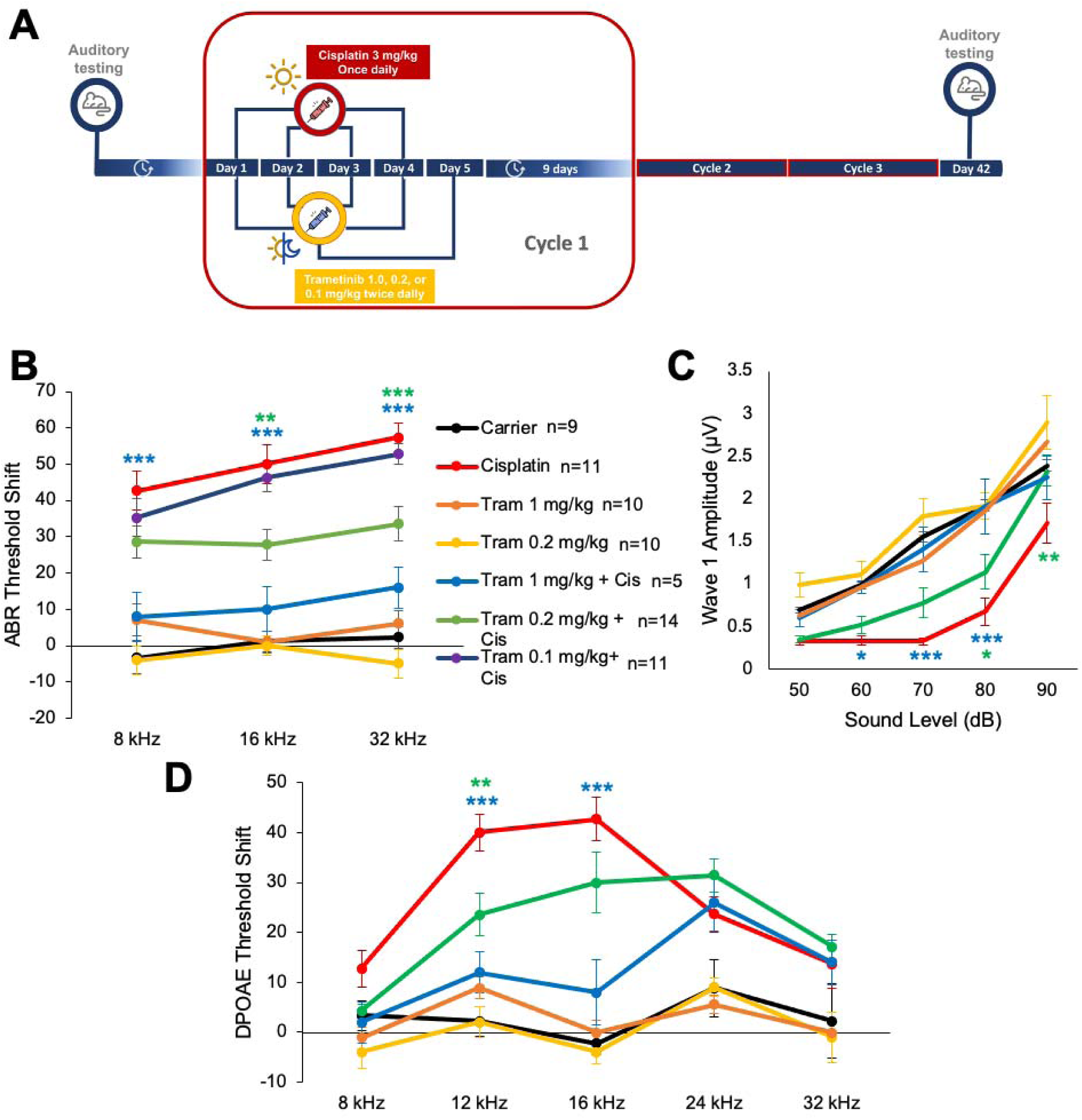
Trametinib protects adult mice from cisplatin-induced hearing loss in a clinically relevant mouse model. **(A)** Schedule of administration of dabrafenib and cisplatin in a translational, multi-cycle cisplatin treatment protocol using CBA/CaJ mice. Each cycle consisted of four days of treatment with 3mg/kg cisplatin in the morning and five days of treatment with 1 or 0.2mg/kg trametinib in the morning and evening. A 9-day recovery period followed the 5 days of treatment. This cycle was repeated for a total of 3 times. Auditory testing occurred before treatment began and immediately after cycle 3 (day 42). **(B)** ABR threshold shifts recorded immediately after the completion of cycle 3 (day 42) in protocol shown in (A). **(C)** Amplitudes of ABR wave 1 at 16 kHz from (B). **(D)** DPOAE threshold shifts recorded after the completion of cycle 3 (day 42) in protocol shown in (A). Carrier alone (black), 1mg/kg trametinib alone (orange), 0.2mg/kg trametinib alone (yellow), cisplatin alone (red), 1mg/kg trametinib + cisplatin (blue), 0.2mg/kg trametinib + cisplatin (green) 0.1mg/kg trametinib + cisplatin (purple). Data shown as means ± SEM, *P<0.05, **P<0.01, ***P<0.001 compared to cisplatin alone by two-way ANOVA with Bonferroni post hoc test.

### Trametinib protects from cisplatin-induced outer hair cell loss

Following the post experiment hearing tests, mouse cochleae were collected and dissections were performed and whole mount images were stained with myosin VI to measure the number of OHC’s in each treatment group. Carrier alone treated mice had an average of 65, 64, and 64 OHCs per 160µm in the apical, middle, and basal regions respectively while cisplatin alone treated mice had an average of 56 OHCs at the apical region, 32 at the middle region, and 4 at the basal region. Mice treated with 1mg/kg trametinib alone had the same number of OHCs compared to the carrier alone cohort at all cochlear regions. Mice co-treated with 1mg/kg trametinib or 0.2mg/kg trametinib with cisplatin had significantly more OHCs at the middle and basal regions compared to cisplatin alone treated mice. Mice co-treated with 1mg/kg trametinib and cisplatin had an average of 59, 56, and 43 OHCs per 160µm at the apical, middle and basal regions, respectively, while mice co-treated with 0.2mg/kg trametinib and cisplatin had an average of 55, 50, and 26 OHCs per 160µm at the apical, middle, and basal regions, respectively. 1mg/kg trametinib plus cisplatin treated mice had a statistically significant higher average of 18 OHCs more per 160µm at the basal region compared to mice treated with 0.2mg/kg trametinib and cisplatin (Figure 3A & B).

**Figure 3:**
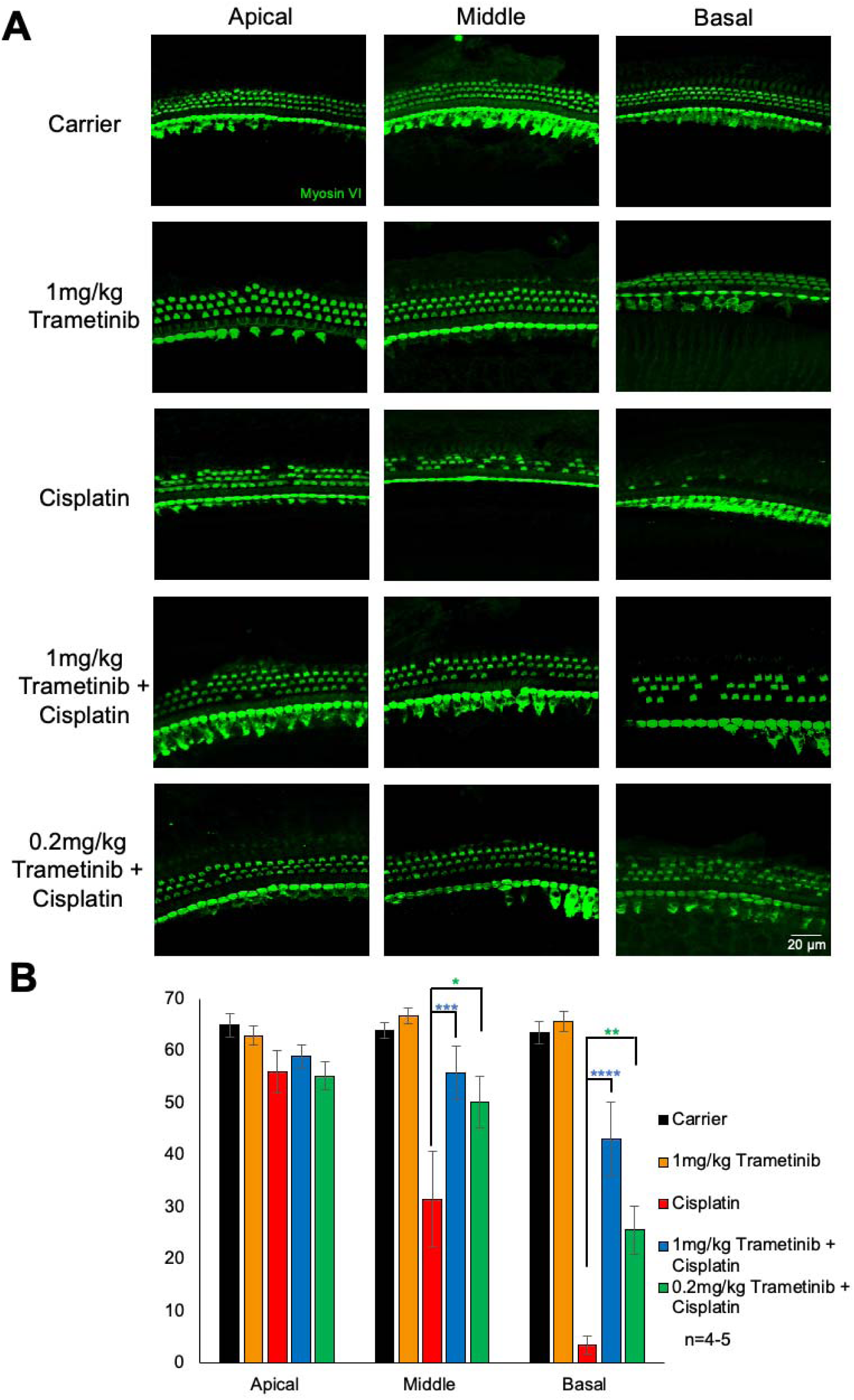
Trametinib protects from cisplatin-induced OHC loss in the multi-cycle cisplatin treatment protocol. **(A)** Representative whole mount cochlear sections stained with myosin VI to visualize hair cells. Treatment groups from top to bottom are as follows: carrier alone, 1mg/kg trametinib alone, cisplatin alone, 1mg/kg trametinib + cisplatin, and 0.2mg/kg trametinib + cisplatin. Apical turn is shown on left, middle turn in the middle, and basal turn on the right. **(B)** Quantification of the number of outer hair cells per 160µm per section for apical turn, middle turn, and basal turn of cochlea. Carrier alone (black), 1mg/kg trametinib alone (orange), cisplatin alone (red), 1mg/kg trametinib + cisplatin (blue), and 0.2mg/kg trametinib + cisplatin (green). Data shown as means ± SEM, *P<0.05, **P<0.01, ***P<0.001, ****P<0.0001 compared to cisplatin alone by two-way ANOVA with Bonferroni post hoc test. n=4-5

### Trametinib confers slight protect from cisplatin-induced weight loss but co-treatment of higher doses of trametinib with cisplatin caused mouse death

Mice were weighed everyday throughout the 42-day treatment protocol shown in Figure 2A. Mice co-treated with both doses of trametinib and cisplatin had significantly less weight loss at days 26, 28, 29, 31, and 39-42 compared to the cisplatin alone treated mice. Cisplatin alone treated mice had a maximum average of 28% weight loss compared to baseline body weight and both trametinib co-treated groups had a maximum average of 22% weight loss throughout the treatment protocol. Trametinib treatment in the absence of cisplatin did not cause any weight loss and mice treated with trametinib alone gradually gained weight throughout the protocol just like carrier alone mice (Figure 4A). The 1mg/kg trametinib co-treated group with cisplatin had significant mouse death with only 36% of mice surviving to the end of treatment protocol while 78% of cisplatin alone treated mice lived to the end of the treatment protocol. 92% of the 0.2mg/kg co-treated mice with cisplatin survived throughout the entire protocol (Figure 4B).

**Figure 4:**
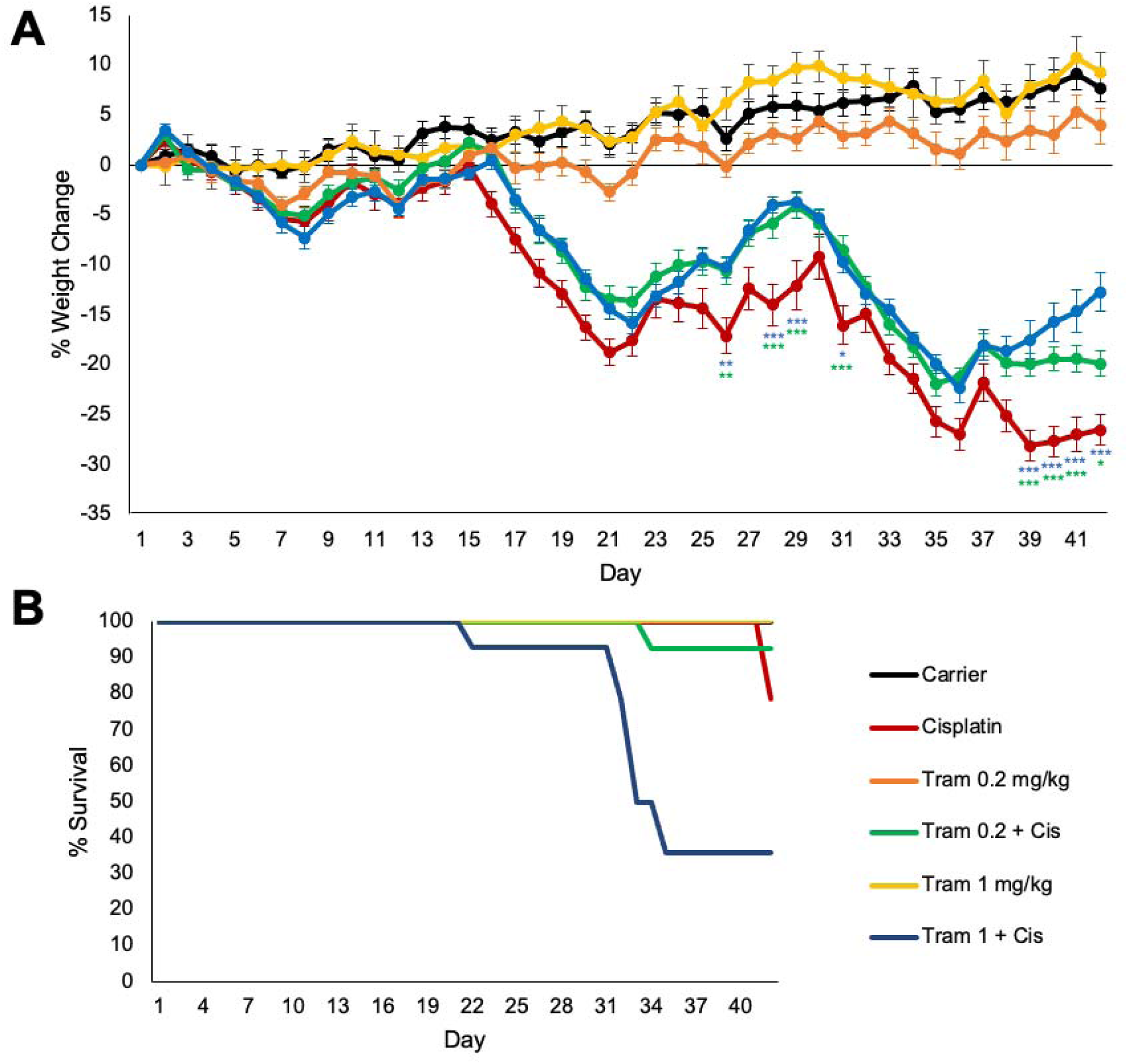
Trametinib confers slight protect from cisplatin-induced weight loss but co-treatment of higher doses of trametinib with cisplatin caused mouse death. **(A)** Weight loss over the 42-day treatment protocol shown in Figure 2A. Data shown as means ± SEM, *P<0.05, **P<0.01, ***P<0.001 compared to cisplatin alone by two-way ANOVA with Bonferroni post hoc test. **(B)** Kaplan-Meier survival curves of mouse cohorts going to day 42 following protocol in Figure 2A. Carrier alone (black), 1mg/kg trametinib alone (orange), 0.2mg/kg trametinib alone (yellow), cisplatin alone (red), 1mg/kg trametinib + cisplatin (blue), and 0.2mg/kg trametinib + cisplatin (green).

### Trametinib protects from noise-induced hearing loss and ribbon synapse loss in FVB mice

To determine whether trametinib protects from noise-induced hearing loss *in vivo,* we utilized a model of NIHL that is performed in FVB mice. Briefly, mice were exposed to 100 dB SPL for 2 hours at 8-16 kHz, which induces permanent threshold shifts in FVB mice, and treatment with 3.15mg/kg trametinib began 24 hours following the noise exposure. Mice were treated twice a day for three days, once in the morning and once in the evening. Hearing tests were performed before and after the treatment protocol to determine how much hearing loss occurred for each treatment group (Figure 5A). As shown in Figure 5B, FVB mice treated with trametinib had significantly lower ABR threshold shifts compared to the noise alone mice. Trametinib treated mice following noise exposure had an average ABR threshold shift decrease of 19 dB at 8 kHz and 18 dB at 16 kHz compared to noise alone mice. Oral administration of trametinib without noise exposure did not induce any hearing loss in the FVB mice (Figure 5B). After post-experimental hearing tests were performed, the cochleae of the mice were harvested and the organ of Corti was dissected to measure ribbon synapse degeneration that is observed following this noise exposure protocol. Outer hair cell loss is not observed in this mouse model of permanent hearing loss but synaptic damage commonly occurs. Myosin VI stained for hair cells and ctbp2 stained the presynaptic ribbon synapses. Example whole mount cochlear images are shown in Figure 5C. The number of ctbp2 puncta per IHC at the 16 kHz region were quantified and trametinib treated mice following noise exposure have significantly more ctbp2 puncta per IHC compared to noise alone mice. Noise + trametinib treated mice have an average of 8.7 ± 0.4 ctbp2 puncta per IHC and noise alone mice have an average of 6.4 ± 0.6 (Figure 5D). Trametinib alone treated mice have an average of 13.4 ± 0.7 ctbp2 puncta per IHC which is comparable to carrier alone treated FVB mice as shown in previous publications [50].

**Figure 5:**
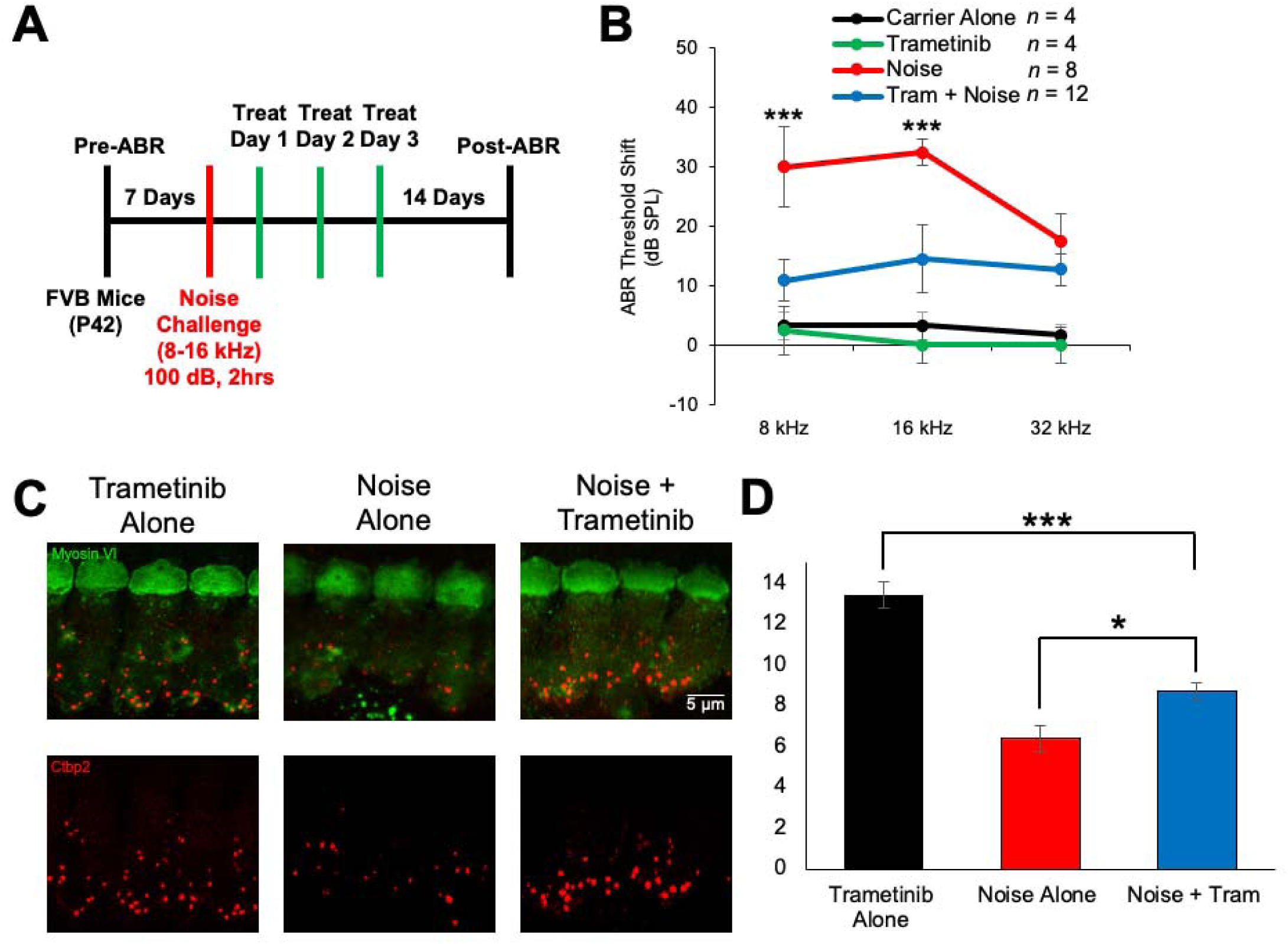
Trametinib protects from noise-induced hearing loss and ribbon synapse loss following noise exposure. **(A)** Noise exposure and treatment protocol. ABR pre-hearing tests were performed and then mice were exposed to 100 dB SPL noise for 2 hours. Starting 24 hours after noise exposure, mice were treated with 3.15mg/kg trametinib twice a day for 3 total days, once in the morning and once at night. 14 days after noise exposure, ABR hearing tests were performed again to determine the hearing loss for each mouse. **(B)** ABR threshold shifts from the treatment protocol shown in (A). Carrier alone (black), trametinib alone (green) noise alone (red), and trametinib + noise (blue). Data shown as means ± SEM, ***P<0.001 compared to cisplatin alone by two-way ANOVA with Bonferroni post hoc test. **(C)** Representative confocal images of whole mount cochlear sections stained with myosin VI (green) and ctbp2 (red). **(D)** Quantification of the average number of ctbp2 puncta per IHC for each treatment group. Trametinib alone (black), noise alone (red), and noise + trametinib (blue). Data shown as means ± SEM, *P<0.05, ***P<0.001 with all groups compared to one another by one-way ANOVA with Bonferroni post hoc test.

## Discussion

This study demonstrates that pharmacological inhibition of MEK1/2 protects from both cisplatin and noise-induced hearing loss. Trametinib prevents MAPK activity by reducing pERK following cisplatin treatment in the HEI-OC1 cell line, as expected due to the selectivity of the drug for MEK1/2 [40]. Trametinib protects mice from cisplatin ototoxicity, as measured by ABR, DPOAE, and outer hair cell counts, while not interfering with cisplatin’s tumor killing efficacy in cell lines. This demonstrates that the protection observed is not through inactivation of cisplatin but through inhibition of the MEK1/2. However, when 1mg/kg of trametinib was co-administered to mice with cisplatin in the multicycle, cisplatin protocol, significant mouse death occurred. A lower dose of 0.2mg/kg trametinib did not cause significant mouse death but still offered protection from hearing loss. Lastly, trametinib did protect from NIHL in FVB/NJ mice while conferring protection from synaptic damage.

Our laboratory has now shown that inhibiting the three main kinases of the MAPK pathway protects from hearing loss. Previous studies have demonstrated that BRAF inhibition protects from both cisplatin and noise-induced hearing loss [24–26]. ERK1/2 inhibition protects from NIHL and now we show that MEK1/2 inhibition protects from both cisplatin and noise-induced hearing loss [27]. Pharmacological inhibition of all three kinases protects from hearing loss in multiple different mouse models. This confirms that the canonical MAPK pathway is leading to hearing loss and that BRAF phosphorylates MEK1/2, which in return phosphorylates ERK1/2 and then cellular pathways downstream of ERK1/2 are activated that contribute to hearing loss (Figure 1A). Very similar hearing protection occurs with inhibition of these three kinases which also shows that these proteins are working through each other and not through other pathways [24–27]. This confirms that the whole MAPK pathway is contributing to hearing loss and not one single kinase in it that could be working through some unknown target.

Interestingly, trametinib was shown to be toxic to mice when combined with cisplatin at a dose of 1mg/kg administered twice a day. This dose is over 4 times more than the human equivalent dose that patients are given for the treatment of melanoma, non-small cell lung cancer, anaplastic thyroid cancer, solid tumors, and glioma [34,51]. This dose was shown to offer significant protection from cisplatin-ototoxicity but only 36% mouse survival was observed when combined with cisplatin. The dose of 0.2 mg/kg given twice a day is approximately the human equivalent dose that is given to patients and no difference in mouse survival was observed compared to the cisplatin alone treated mice [51]. The dose of 1mg/kg was toxic to the mice, most likely because the dose was much higher than what has been demonstrated to be safe. Even though trametinib has been well tolerated in patients, it appears to be toxic when combined with cisplatin at higher doses, which is why dabrafenib is such a promising drug to repurpose for the protection of cisplatin and noise-induced hearing loss. Dabrafenib offers the same protection as MEK1/2 inhibition; however, the toxicity profile is very favorable, and we have demonstrated that dabrafenib does not cause any known systemic toxicity when co-administered with cisplatin and even protects mice from weight loss associated with cisplatin treatment [24]. The therapeutic window for trametinib to protect from cisplatin ototoxicity is less than 5 and we have previously shown that dabrafenib’s therapeutic window is greater than 25 in the multicycle, cisplatin mouse model [24]. Trametinib does confer significant protection from cisplatin ototoxicity; however, dabrafenib is a much more promising otoprotective agent because of its safety profile and high efficacy of protection from hearing loss in preclinical studies.

Trametinib treatment protected from noise-induced synaptic damage as demonstrated by more ctbp2 puncta in the treated group compared to the noise-alone group. Dabrafenib (BRAF inhibitor), tizaterkib (ERK1/2 inhibitor), and now trametinib (MEK1/2 inhibitor) have all demonstrated significant protection from noise-induced synaptopathy [24–27]. With three inhibitors of three separate kinases of the MAPK pathway offering protection from synaptopathy, this suggests that one of the major mechanisms of protection from hearing loss with MAPK inhibition is through prevention of synaptic damage [52–54]. Overactivation of the MAPK pathway is leading to synaptic damage following noise exposure and pharmacological inhibition protects this from occurring in mice. Exactly how MAPK inhibition is protecting from synaptopathy is not yet known; however, we have demonstrated that ERK1/2 inhibition does prevent immune cell infiltration following noise exposure which contributes to the hearing loss and possibly synaptopathy [27]. MAPK activation has been shown to induce a pro-inflammatory environment and we have demonstrated that MAPK inhibition can reduce this pro-inflammatory response following noise exposure [27,33,55–58]. Additionally, MAPK activation has been demonstrated to occur in the supporting cells of the inner ear following cisplatin administration and noise exposure, specifically in the Deiters’ and inner phalangeal cells, and these supporting cells help clear glutamate from the synapses between hair cells and the nerve fibers [24–26]. Lowering MAPK activity in the supporting cells could be helping with supporting cell health which would allow for proper functioning of these cell types, such as clearing out glutamate from the synapses [59–61]. Glutamate excitotoxicity is known to occur following both cisplatin and noise exposure and this leads to synaptic dysfunction and cell death [62,63]. Preventing supporting cell dysfunction and death could help clear glutamate out of the synapses which would reduce the synaptopathy that occurs from these ototoxic insults [59,61]. This is a very interesting mechanism that MAPK inhibition is protecting from hearing loss through, and further research is needed to elucidate exactly how this is occurring.

In summary, trametinib protects from both cisplatin and noise-induced hearing loss. Trametinib treated mice had lower ABR and DPOAE threshold shifts compared to noise and cisplatin alone treated mice while also having significant protection from synaptopathy and outer hair cell loss. BRAF, MEK1/2, and ERK1/2 inhibition all protect from NIHL while BRAF and MEK1/2 inhibition protect from cisplatin ototoxicity [24–27]. Trametinib was shown to be toxic to mice at high doses when co-treated with cisplatin, but lower doses did not show any obvious deleterious side effects. This study shows that activation of the canonical MAPK pathway is leading to hearing loss and inhibiting any of these kinases leads to hearing protection. This study also demonstrates that dabrafenib is the most promising MAPK inhibitor to be repurposed for hearing loss prevention because it has a very favorable safety profile and confers significant protection from both insults of hearing loss [24–26].

## Materials and Methods

### Study Approval

All animal experiments included in this study were approved by Creighton University’s Institutional Animal Care and Use Committee (IACUC) in accordance with policies established by the Animal Welfare Act (AWA) and Public Health Service (PHS).

### Mouse Models

For the cisplatin studies, 8-week-old CBA/CAJ mice were purchased from Jackson Laboratory (Bar Harbor, Maine, USA). Mice were then allowed to acclimate to the Animal Resource Facilities (ARF) at Creighton University before any experimental procedures began. For the noise studies, FVB/NJ breeding mice were purchased from Jackson Laboratory, bred in the ARF at Creighton University, and experiments were performed on the offspring of the breeders when mice were 8-10 weeks old. For pre- and post-experimental hearing tests, mice were anesthetized by Avertin (2,2,2-tribromoethanol) via intraperitoneal injection at a dose of 500 mg/kg, and complete anesthetization was determined via toe pinch. For all experiments, mice were randomly assigned to experimental groups with equal numbers of male and females.

### HEI-OC1 Cell Line and Collection of Cell Lysates

The HEI-OC1 cell line was maintained with DMEM 1× with glucose (4.5 g/liter), l-glutamine, and sodium pyruvate (12430-054, Gibco) with 10% FBS and ampicillin (50 μg/ml) added to the media. Cells were cultured under the conditions of 33°C and 10% CO_2_ while being passaged with 0.05% trypsin/EDTA. For the collection of cell lysates, HEI-OC1 cells were grown in T-75 flasks and allowed to grow to 80-90% confluency. 6 separate flasks of cells were used with each flask being its own treatment group. The 6 treatment groups were medium alone, 1μM trametinib alone, 50μM cisplatin alone, 50μM cisplatin + 0.01μM trametinib, 50μM cisplatin + 0.1μM trametinib, and 50μM cisplatin + 1μM trametinib. HEI-OC1 cells were pre-treated with trametinib for 1 hour which was followed with cisplatin treatment for 1 hour. After one hour of cisplatin treatment, cell lysates were collected by dumping the media, putting cold DPBS in each flask, and scrapping the bottom of the flask with a cell scrapper to collect the cells in the DPBS. This was performed for a total of 4 separate times for each experimental group. Cells were then centrifuged at 2000 RPM for 5 minutes at 4°C to form the cell pellet. Lysis buffer was then added to the cell pellet, cells were resuspended in the lysis buffer, and centrifuged for 20 minutes at 16,000g at 4°C. The supernatant was then collected and put in −80°C for storage before western blots were performed.

### Cancer Cell Lines and Cell-Titer Glo Assay

For tumor cell lines viability experiments, two neuroblastoma (SK-N-AS and SH-SY5Y) and a non-small cell lung carcinoma cell line (A549) were utilized. 9600 cells per well were plated in 6-replicates in 96 well plates and allowed to attach overnight at 37°C in 5% CO_2_. The following day, the tumor cell lines were pretreated with a range of 30µM-4.57nM of trametinib with 1/3 serial dilutions starting at 30µM to 4.57nM. The cells were then treated with cisplatin and incubated for 48 hours. The cisplatin concentration was dependent on the cell line and the IC_50_ of cisplatin for the respective cell line was used as previously determined. The A549 cell line was treated with 25µM cisplatin, SK-N-AS was 6.5µM, and SH-SY5Y was 2µM. Cell viability was then measured using the Cell Titer-Glo (Promega) assay. Medium alone-, cisplatin alone-, and drug alone were used as controls and the percent viability was calculated as the viability compared to the medium alone treated cells. Drug plus cisplatin treated cells were compared to cisplatin alone treated cells to determine whether trametinib interfered with cisplatin’s tumor killing ability.

### Immunoblotting

Cell lysates that were stored in the −80°C were thawed, and protein concentrations were determined with the BCA protein assay kit (23235, Thermo Fisher Scientific). Thirty micrograms of total cell lysate were loaded on 10% SDS–polyacrylamide gel electrophoresis gels. After running the gel and transferring to a nitrocellulose membrane, the following antibodies were used for immunoblotting analysis: anti-ERK1/2 (4695, Cell Signaling Technologies, 1:1000) anti-pERK1/2 (Thr^202^/Tyr^204^, 9101S, Cell Signaling Technologies, 1:1000) and anti-GAPDH (ab181602, abcam, 1:5000). Anti-rabbit (P0448) secondary antibody was purchased from Dako Laboratories and diluted 1:4000. National Institutes of Health (NIH) ImageJ software was used to quantify the band intensities and recorded as a ratio to the loading control GAPDH.

### Multi-cycle cisplatin treatment model

Pre-experimental ABR were performed on 9-week-old CBA/CaJ mice with DPAOE performed when mice were 10 weeks old. Once mice were 12 weeks old, the 6-week cisplatin and trametinib treatment regimen began. Trametinib (1, 0.2, or 0.1mg/kg) was administered via oral gavage 1 hour before 3 mg/kg cisplatin was administered to mice via intraperitoneal injection in the morning. Mice were then treated with trametinib or carrier again in the evening. Mice were treated with cisplatin once a day for 4 days and trametinib twice a day for 5 days with a 9-day recovery period in which no drugs were administered to the mice. This cycle was repeated two more times for a total of 3 cycles. Mice were treated with 3 mg/kg cisplatin for a total of 12 days (4 days per cycle with 3 cycles) and trametinib for a total of 15 days (5 days per cycle with 3 cycles). Immediately after the completion of cycle 3 (42 days after the first cisplatin injection), post-experimental ABR were performed with DPOAE performed one week after ABR. Cochleae were when harvested and put in 4% PFA. Cisplatin treated mice were injected by subcutaneous injection twice a day with 1 mL of warm saline to ameliorate dehydration. This continued until body weight started to recover. Mouse weight was analyzed everyday throughout the 42-day treatment protocol. The cages of cisplatin-treated mice were placed on heating pads throughout the duration of the experiment until mice began to recover after the 3rd treatment cycle of the protocol. Food pellets dipped in DietGel Boost® were placed on the cage floor of cisplatin-treated mice. The investigators and veterinary staff carefully monitored for changes in overall health and activity that may have resulted from cisplatin treatment.

### Auditory Brainstem Response

ABR waveforms in anesthetized mice were recorded in a sound booth by using subdermal needles positioned in the skull, below the pinna, and at the base of the tail, and the responses were fed into a low-impedance Medusa digital biological amplifier system (RA4L; TDT; 20-dB gain). Mice were anesthetized by 500 mg/kg Avertin (2,2,2-Tribromoethanal, T4, 840-2; Sigma-Aldrich) with full anesthesia determined via toe pinch. At the tested frequencies (8, 16, and 32 kHz), the stimulus intensity was reduced in 10-dB steps from 90 to 10 dB to determine the hearing threshold. ABR waveforms were averaged in response to 500 tone bursts with the recorded signals filtered by a band-pass filter from 300 Hz to 3 kHz. ABR threshold was determined by the presence of at least 3 of the 5 waveform peaks. For noise exposure experiments, baseline ABR recordings were performed when mice were 7-8 weeks old and post experimental recordings were performed 14 days after noise exposure. For cisplatin experiments, ABR recordings were performed when mice were 9 weeks old and post experimental recordings were performed after the completion of the 42-day treatment protocol when mice were 18 weeks old. All beginning threshold values were between 10 and 40 dB at all tested frequencies. All thresholds were determined independently by two-three experimenters for each mouse who were blind to the treatment the mice received. Threshold shifts were calculated by subtracting the pre-noise exposure recording from the post-noise exposure recording. ABR wave one amplitudes were measured as the difference between the peak of wave 1 and the noise floor of the ABR trace.

### Distortion Product Otoacoustic Emission

Distortion product otoacoustic emissions were recorded in a sound booth while mice were anesthetized. Mice were anesthetized by 500 mg/kg Avertin (2,2,2-Tribromoethanal, T4, 840-2; Sigma-Aldrich) with full anesthesia determined via toe pinch. DPOAE measurements were recorded using the TDT RZ6 processor and BioSigTZ software. The ER10B+ microphone system was inserted into the ear canal in way that allowed for the path to the tympanic membrane to be unobstructed. DPOAE measurements occurred at 8, 12, 16, 24, and 32 kHz with an f2/f1 ratio of 1.2. Tone 1 was *.909 of the center frequency and tone 2 was *1.09 of the center frequency. DPOAE data was recorded every 20.97 milliseconds and averaged 512 times at each intensity level and frequency. At each tested frequency, the stimulus intensity was reduced in 10 dB steps starting at 90 dB and ending at 10 dB. DPOAE threshold was determined by the presence an emission above the noise floor. For noise exposure experiments, baseline DPOAE recordings were performed when mice were 7-8 weeks old and post experimental recordings occurred after 14 days following noise exposure. For cisplatin experiments, ABR recordings were performed when mice were 10 weeks old and post experimental recordings were performed after the completion of the 42-day treatment protocol when mice were 19 weeks old. Threshold shifts were calculated by subtracting the pre-noise exposure recording from the post-noise exposure recording.

### Noise Exposure

Mice were placed in individual cages in a custom-made wire container. A System RZ6 (TDT) equipment produced the sound stimulus which was amplified using a 75-A power amplifier (Crown). A JBL speaker delivered the sound to the mice in the individual chambers. The sound pressure level was calibrated using an NSRT-mk3 (convergence instruments) microphone and all chambers were within 0.5 dB of each other to ensure equal noise exposure. Mice were exposed to 100 dB SPL noise for 2 hours with an octave band noise of 8 to 16 kHz.

### Tissue preparation, immunofluorescence, and OHCs counts

For OHC counts, cochleae from adult mice were prepared and examined as described previously [24,25,64]. Cochleae were placed in 4% PFA following harvesting. Cochlear samples were stained with anti-myosin VI (1:400; 25-6791, Proteus Bioscience) with the secondary antibody purchased from Invitrogen coupled to anti-rabbit Alexa Fluor 488 (1:400; A11034). All images were acquired with a confocal microscope (LSM 700 or 710, Zeiss). Outer hair cell counts were determined by the total amount of outer hair cells in a 160 µm region. Counts were determined for the 8, 16, and 32 kHz regions. Cochleae from each experimental group were randomly selected to be imaged for outer hair cell counts.

For ctbp2 counts, cochleae were harvested and prepared the same as cochleae for OHC counts. Organs of Corti were micro dissected and co-stained with anti-Ctbp2 (1:800; 612044, BD Transduction) and myosin VI (1:400; 25-6791, Proteus Biosciences). Goat anti-rabbit Alexa Fluor 488 (1:400; A11034) and goat anti-mouse Alexa Fluor 647 (1:800; A32728) were purchased from Invitrogen as the secondary antibodies. Confocal Imaging was performed using a Zeiss 700 upright scanning confocal microscope with images taken with the 63x objective lens. Final images were achieved by taking a z stack image of the organ of Corti and processed through the ZenBlack program. The number of Ctbp2 puncta were counted per IHC with a total of 12-14 IHCs analyzed at the 16 kHz region. The total amount of Ctbp2 puncta were divided by the total amount of IHCs in the 16 kHz region to determine the number of Ctbp2 puncta per IHC.

### Statistical Analysis

Statistics was performed using Prism (GraphPad Software). Two-way analysis of variance (ANOVA) or One-way ANOVA with Bonferroni post hoc test was used to determine mean difference and statistical significance. Specific statistical tests used for each experiment are in the figure legends. Statistical significance was determined when P <0.05.

## Data and materials availability

All data needed to evaluate the conclusions in the paper are present in the paper and/or Supplementary Materials. Additional data can be requested from the authors if needed.

## Acknowledgements

We thank Daniel F. Kresock, Kristina Ly, Dr. Christy Howe, Dr. Janee Gelineau-van Waes, Pat Steele, Ann Bryen and the Creighton University ARF staff for assistance with the mouse studies. The research was funded by the National Institutes of Health NIDCD grant 1R01DC018850, American Hearing Research Foundation 2020 grant to T. Teitz, and National Institutes of Health NIDCD grant 1F32DC020102 to M. Ingersoll. This investigation was conducted in facilities constructed with support from Research Facilities Improvement Program (G20 RR024001-01) from the National Center for Research Resources, NIH. The research was partially conducted at the Auditory and Vestibular Technology Core (AVT) at Creighton University, Omaha, NE (RRID:SCR_023866). This facility is supported by the Creighton University School of Medicine and grants GM103427 and GM139762 from the National Institute of General Medical Science (NIGMS), a component of the National Institutes of Health (NIH). IBIF was constructed with support from grants from the National Center for Research Resources (RR016469) and the NIGMS (GM103427). This investigation is solely the responsibility of the authors and does not necessarily represent the official views of the National Center for Research resources, NIGMS or NIH.

## Author Contributions

T.T. and M.A.I. conceived the project. R.D.L. and M.A.I. performed the HEI-OC1 studies and western blots. R.D.L. performed the *in vitro* cisplatin interference experiments. R.D.L., M.A.I., and R.G.K. performed the mouse treatments for the cisplatin and noise studies. R.D.L., M.A.I., and R.G.K. performed ABRs and DPOAEs. R.D.L. performed the cochlear dissections and staining for the OHC counts and Ctbp2 counts. R.D.L. and T.T. wrote the manuscript with input from the other authors.

## Conflicts of Interest

T.T. is an inventor on US patent no. 11,433,073 and US serial no 17/736,330 filed for the use of trametinib for hearing protection, and patent application 62/500,677;WO2018204226, for the use of dabrafenib for hearing protection, and is a cofounder of Ting Therapeutics LLC.

